# Force-based three-dimensional model predicts mechanical drivers of cell sorting

**DOI:** 10.1101/308718

**Authors:** Christopher Revell, Raphael Blumenfeld, Kevin Chalut

**Author notes:** Corresponding author Email address (Kevin Chalut).

## Abstract

Many biological processes, including tissue morphogenesis, are driven by mechanical sorting. However, the primary mechanical drivers of cell sorting remain controversial, in part because there remains a lack of appropriate threedimensional computational methods to probe the mechanical interactions that drive sorting. To address this important issue, we developed a three-dimensional, local force-based simulation method to enable such investigation into the sorting mechanisms of multicellular aggregates. Our method utilises the subcellular element method, in which cells are modeled as collections of locally-interacting force-bearing elements, accommodating cell growth and cell division. We define two different types of intracellular elements, assigning different attributes to boundary elements to model a cell cortex, which mediates the interfacial interaction between different cells. By tuning interfacial adhesion and tension in each cell cortex, we can control and predict the degree of sorting in cellular aggregates. The method is validated by comparing the interface areas of simulated cell doublets to experimental data and to previous theoretical work. We then define numerical measures of sorting and investigate the effects of mechanical parameters on the extent of sorting in multicellular aggregates. Using this method, we find that a minimum adhesion is required for differential interfacial tension to produce inside-out sorting of two cell types with different mechanical phenotypes. We predict the value of the minimum adhesion, which is in excellent agreement with observations in several developmental systems. We also predict the level of tension asymmetry needed for robust sorting. The generality and flexibility of the method make it applicable to tissue self-organization in a myriad of biological processes, such as tumorigenesis and embryogenesis.

## 1. Introduction

Self-organization is a widely-studied feature of non-equilibrium systems across a great variety of fields [1, 2, 3]. In Biology, self-organized processes range in length-and time-scales from protein folding [4] to the dramatic murmurations of starlings [5]. Generically, these systems exhibit emergence of macroscopic order from disordered states by the action of simple, local, inter-component interactions. Here we focus on one such a process of self-organization - cell sorting. Cell sorting in a multicellular aggregate (MCA) is critical for normal development of an embryo, and is also at the heart of pathologies such as metastastic growth of tumors.

The spontaneous separation of mixtures of embryonic cells has long been thought to be driven by cells’ differing “affinity” for one another [6]. Two primary drivers have been proposed to facilitate differential affinity: differential adhesion and differential interfacial tension.

The differential adhesion hypothesis is a widely studied driver of cell sorting [7, 8, 9]. It asserts that, in an MCA with multiple cell types, those with strong mutual adhesion adhere together better than cells with weaker mutual adhesion. The latter are ‘sorted’ to the outside of the MCA, resulting in cellular segregation. Although the differential adhesion hypothesis has been applied to a number of systems [10], it has been challenged in the light of observations that sorting occurs even when two cell types adhere to one another equally well [11], and that contractile forces are more responsible than adhesion for tissue mor-phogenesis in a number of systems [12, 13].

The idea that sorting in many biological processes is driven primarily by contractile forces is at the heart of the differential interfacial tension hypothesis [14], the basic postulates of which are: (i) cell surface membranes exhibit local tension variation [15]; (ii) a cell surface tension is highest when in contact with the external medium and lowest when in contact with cells of the same type [16]. The variation in surface tension at cell interfaces leads to variation in cell interface area, and hence to mutual affinity [Figure 2a]. According to the differential interfacial tension hypothesis, variations in mutual affinity, driven by differences in surface tension, are primarily responsible for self-organization in a cell aggregate. The reduction in tension at cell interfaces is proposed to occur by the exclusion of myosin from the actomyosin cortex at the interface. Such exclusion is caused by the intra-membrane proteins that mediate adhesion between the cells [13]. This passive mechanism allows morphogenesis to proceed without active movements by the cells involved and, moreoever, provides the local coordination between cells needed for complex global morphogenesis [17].

The importance of intercellular differential interfacial tension for tissue morphogenesis and maintenance has been demonstrated in a number of systems. For example, it is responsible for boundary maintenance in the the dorsoventral compartment boundary in the *Drosophila* wing imaginal disc [18, 19]. Much of this body of work has focussed on 2D epithelial systems, often maintaining boundaries rather than forming boundaries from a mixed aggregate [20]. However, further evidence of the importance of differential interfacial tension comes from experimental work on 3D aggregates, suggesting that local variation in cortical tension is responsible for internalizing the first set of internal cells in the mouse morula [21]. Moreoever, reduction in interfacial tension has also been shown to drive morula compaction [22] and allocation of cells to the inner cell mass of the embryo [23].

In order to investigate in detail the effect of differential interfacial tension on 3D MCAs, we constructed a computational model based on the subcellular element method (SCEM) [?]. To validate the method, we compared its predictions to theoretical models of differential interfacial tension in cell doublets [13] [Figure 2c]. We used our model to simulate sorting in 3D MCAs of two cell types. Using our model, we are able to predict the minimum adhesion magnitude required for tension-driven sorting to occur. We show that, given this minimum adhesion, interfacial tension asymmetries drive sorting in multi-cellular aggregates. We also show that fairly significant differential adhesion is necessary for cell sorting. Ultimately, we also quantitatively predict the amount of tension asymmetry necessary for sorting.

## 2. Subcellulaг Element Method (SCEM)

A number of techniques have been developed to model cellular systems [24, 25, 26, 27], including explorations of differential interfacial tension using the vertex method in two dimensional epithelia [19, 28]. However, many of these methods rely on a global Hamiltonian, which is a strong, abstract assumption for modelling biological systems that do not necessarily conserve energy. Instead, we set out to develop a computational model that instead assumes dynamics driven by local forces. We required a method that allows integration of subcellular-scale mechanics, including stiffness, tension, and adhesion, with 3D tissue-scale phenomena. Therefore, we chose to implement SCEM.

SCEM is a method for multi-scale 3D simulation that can simultaneously model intra-and inter-cellular dynamics, as well as the development of multicel-lular tissues. It allows us to study the effects of cell shape changes and complex inter-and intra-cell features on tissue-scale dynamics [29]. The method treats each cell as a collection of elements, interacting via nearest-neighbor forces [Figure 1]. Nearest-neighbor elements of different cells also interact, facilitating intercellular interactions by the same mechanism. All the inter-element interactions are assumed to be via a Morse potential [30], which is generically repulsive at short distances with a minimum at some equilibrium distance. Different element types, however, have different potential parameters. For the purposes of this work, we modified and extended the SCEM simulation code significantly from its original version [31].

**Figure 1:**
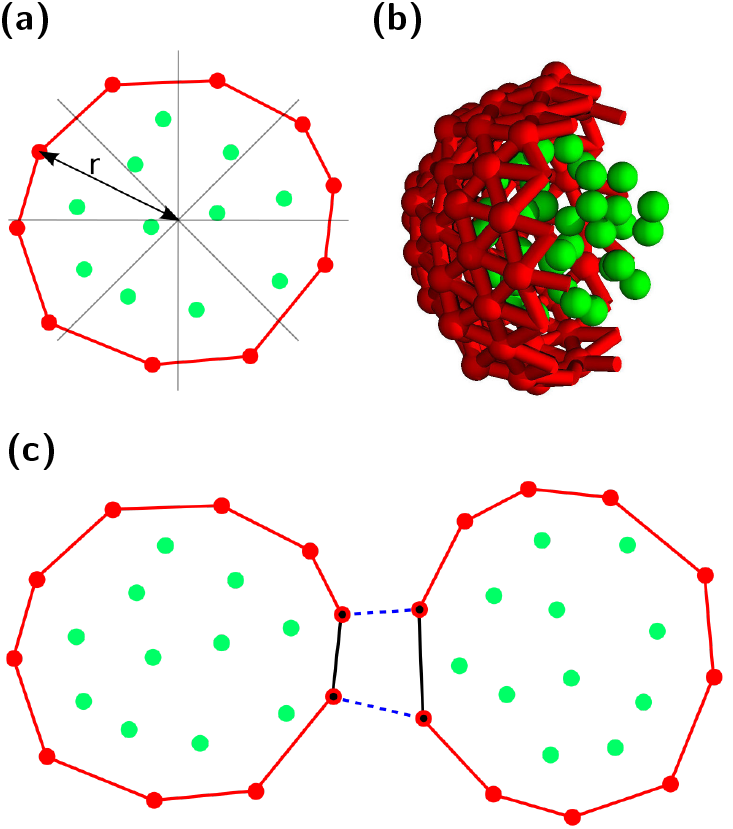
(a), Algorithm for allocation of cortex elements, 2D representation. Cell is divided into π/4 radian bins. Elements with a radius greater than 80% of the maximum radius in their bin are allocated cortex type. (b), Cutaway of one SCEM cell showing all cytoplasm elements and part of the cortical tension network as defined by a Delaunay triangulation. (c), Algorithm for implementing differential interfacial tension between two cells. Solid lines indicate cortical tension forces; dotted lines indicate inter-cell adhesive interactions. Elements that share adhesive interactions with another cell are labelled according to the type of cell, shown with a black dot. An interface is defined as a region in which cortex elements have the same label. The tension is altered for any cortex interaction between two cortex elements with the same label, shown by solid black lines. Cytoplasm elements are green and cortex elements are red throughout.

An essential modification of SCEM was an addition of a contractile cortex to each cell, the properties of which are different from the main bulk of the cell. This allowed us to control the cell cortical tensions. The modification involved first identifying the surface elements of the cell. This was done by dividing the cell into 32 sectors of equal steradians, emanating from its center of mass, and defining as cortex elements all sector elements within 20% of the radius distance to the furthest element in each sector, as illustrated in Fig. 1a.

To ensure that the interaction between cortex elements is tangential to the cell surface, we performed a Delaunay triangulation [32, 33] over each cell’s cortex elements, which makes the cell a polyhedron covered in triangular facets. By its nature, this triangulation favors roughly equilateral triangles, which produces a relatively smooth cell surface [Fig. 1b]. Two cortex elements sharing such a triangle edge are neighbors and they interact along the edge via a tensile force. Thus defined, the cortex forces are guaranteed to span the full cell surface, be tangential to it, prevent elements detaching from the cell, and enable the modelling of intercellular mechanical interactions.

With our computational method, the cortex tension can vary locally across the cell surface in response to contact with surfaces of other cells. Such variation is achieved by changing the magnitude of intra-cell tension forces between cortex elements that lie at a cell interface. Specifically, we define a cell interface by first identifying the cortex elements that share an adhesive inter-cell interaction with a cortex element of a different cell. These elements are then labeled according to whether the other cell is of the same or different fate. An interface between the two cells is then defined as a region containing only cortex elements of the same label, and any cortical tension force acting between two cortex elements with the same label can be altered [Fig. 1c]. The magnitude of intercellular tension asymmetry can be specified differently depending on cell type and whether the interface is between like or unlike cells.

Changing the local interaction between cell cortex elements affects not only the local cortical tension but also the distance between the elements and, there fore, the local element density. Since intercellular adhesion is mediated by these elements, an increase in element density, for example, increases the local adhesion strength between neighboring cells. To compensate for this, we normalize the adhesion magnitude by the local element density [Appendix A].

## 3. Simulation of cell doublets

The simplest experimental system to define the mechanical phenotype of cells and their interfaces is a cell doublet, for which in-vitro experiments and theoretical models exist. This makes this system a suitable test case to validate our method. We expect sorting to be driven by changes in relative affinity, reflected by changes in equilibrium interfacial contact area (or, analogously, contact angle) between cells. This interfacial contact area depends upon the adhesion magnitude between cells (*ω*) the cortical tension of the cells (*γ_m_*), and the interfacial tension (*γ*_*c*_).

Assuming an symmetric cell doublet is in mechanical equilibrium, the dependence of the contact angle between two cells on their surface tensions can be deduced using linear force balance at the vertex [13] [Figure 2c]. Further assuming, based on experimental observations [34, 35, 13], that the effect of *ω* is negligible in the high-tension limit, one can derive a relationship between *γ_m_, γ*_*c*_, and the doublet contact angle *θ* [Eq. 1].

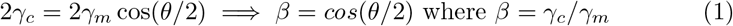

**Figure 2:**
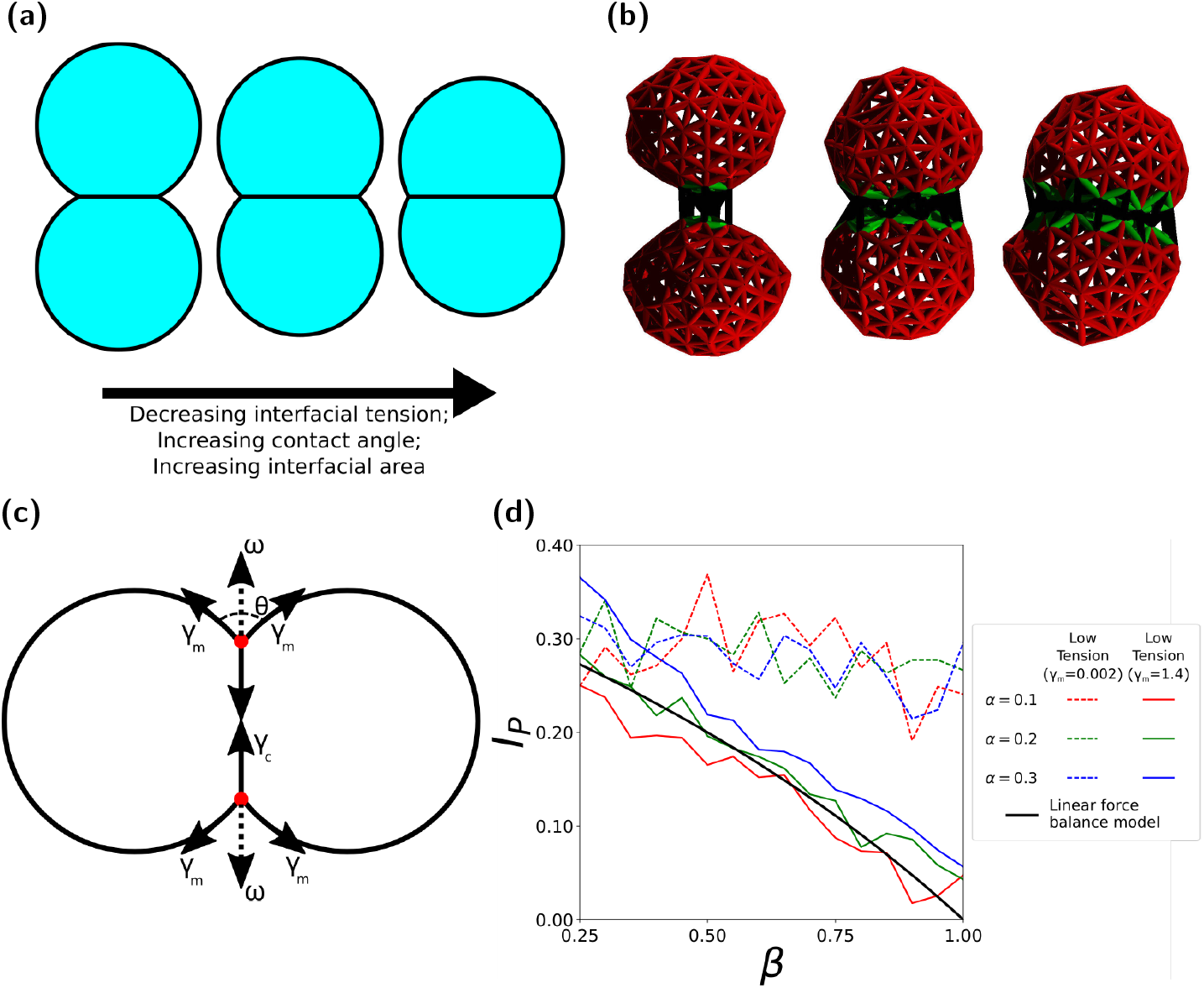
(a), Two dimensional diagram of cell doublets showing the effect of changing interfacial tension on contact angle and interface area. Interfacial tension decreases from left to right, resulting in increased interfacial area, increased contact angle, and consequently increased mutual affinity between cells. (b), Cell doublets visualised by cortical tension force network, demonstrating variation of interface area for interfacial tension factor values *β* = 1.00, 0.75, 0.50. (c), Diagram linear force balance of cell doublet contact area, as presented in [13]. Cylindrical symmetry allows the system to be reduced to a two dimensional force balance at the vertices of the cell interface. The influence of adhesion is reframed as a colinear tension force ω acting to pull the vertices apart. In order to reach equilibrium, the forces pulling the edges apart must balance the forces pulling the edges together. (d), Plots of interface proportion against *β* in high tension and low tension regimes.

To test the effect of system parameters on cell doublets in our simulations, we allow an initial cell to divide into two, forcing both to be of the same type. We then allow the system to reach mechanical equilibrium without growth or division, producing a doublet of identical cells, adhered at a joint interface [Figure 2b].

Experimentally, a straightforward indicator of cell affinity in doublets is the contact angle between the two cells, *θ*. While this measurement is intractable in our simulations, we could measure straightforwardly the cell-cell interface area as the sum the areas of all the Delaunay triangles. Using simple trigonometry, we found the ratio, *I_P_*, of the interfacial contact areas in simulations to the total cell surface area and related it to the contact angles obtained in experiments, *θ*:

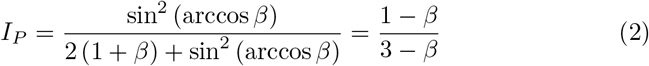

This allows direct comparison of our simulations to linear force balance at cell interfaces [Equation 2] and provides a validation test of our method’s predictions [Figure 2d]. We define an adhesion magnitude *A_M_* for our simulations, as distinguished from *ω* because it reflects the sum of the local adhesion magnitudes between intercellular elements. The validation consisted of simulating cell doublets, from which we obtained measurements of *I_P_* for values of *β* between 0.25 and 1. We used a representative set of *A_M_* and *γ_m_* values, corresponding to low tension and high tension regimes. The resulting *I_P_* values were then compared to the theoretical predictions of linear force balance [Fig. 2d].

We found very good agreement between the simulation results and the theoretical prediction of the force balance model in the high tension regime (solid lines in the figure), with adhesion producing a small constant offset across *β*. The magnitude of this offset increases gradually with adhesion. In contrast, in the low tension regime, adhesion cannot be ignored when considering the linear force balance in a cell doublet. Specifically, we observe that *I_P_* depends only weakly on *β*, confirming the constraints of using linear force balance to predict cell-cell contacts and therefore sorting of different cell types. Most importantly, we conclude that due to the excellent agreement with linear force balance in the high tension regime, our method can be used to predict the mechanical interactions within MCAs.

By simulating a large number of cell doublets at different parameters, we constructed phase diagrams for the behavior of the cell doublet interface area as a function of *A_M_* and *γ_m_* for *β* = 0.5, 0.75, and 1.00 [Figs. 3a-3c]. The phase diagrams indicate that the highest *I_P_* value achieved for any parameter set is approximately 0.32. This value is in good agreement with the theoretical limit for the interface between two hemispheres, which is exactly 1/3. Our doublet simulations also show that, for each value of *β*, the measurement of *I*_*P*_ drops sharply with increasing *γ_m_*, tending to a minimum [Fig. 3f]. The minimum is determined by *A_M_*, with very low adhesions producing little to no interface. Increasing adhesion from zero produces a sharp increase in interface, but beyond a threshold adhesion the increase of interface with adhesion magnitude has a much smaller gradient [Fig. 3d].

**Figure 3:**
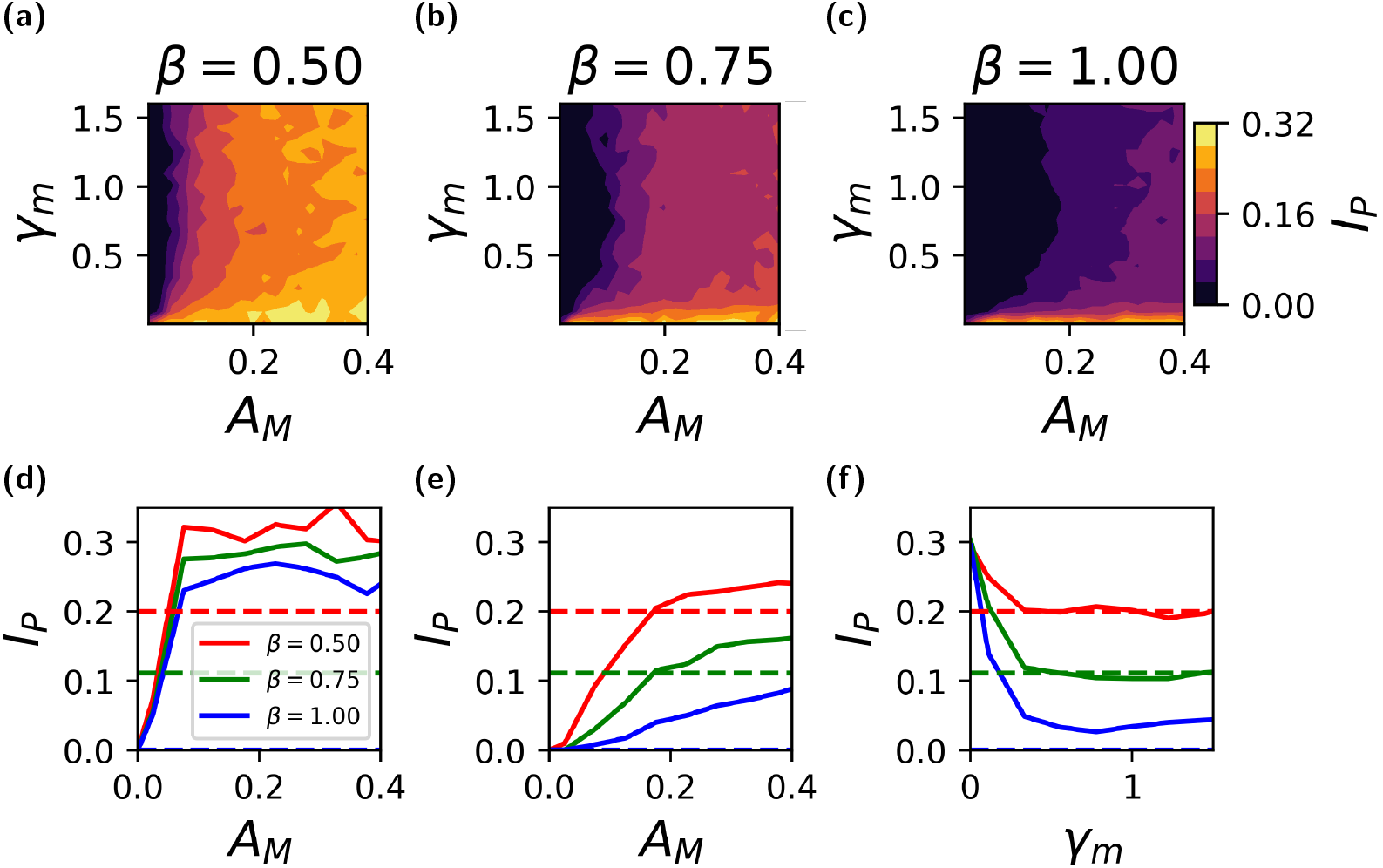
(a-c) Phase space plots showing variation of interface proportion (*I_P_*) in *γ_m_* and *A_m_* space measured in cell doublet simulations. (a), *β* = 0.5. (b), *β* = 0.75. (c), *β* = 1.0. Force balance predicts *I*_*P*_ = 0.20, 0.11, 0.00 respectively. (d-f), Plots of interface area against adhesion and tension magnitude. Dotted lines show predictions made by balancing forces. (d), Interface against adhesion in low tension regime, *γ_m_* = 0.01. (e), Interface against adhesion in high tension regime, *γ_m_* = 1.2. (f), Interface against tension magnitude at *A_M_* = 0.2.

The doublet simulation results are even clearer if we take slices through the phase diagram [Figs. 3d-3f]. As expected, we find that, for both low and high tensions, little to no interface is produced at low adhesion values. However, for both low and high tension the interface increases approximately linearly with increasing adhesion. In the low tension regime [Fig. 3d], interfacial contact area increases sharply with adhesion until a value of approximately *A_M_* = 0.1, after which the interfacial contact area plateaus. The value at which the interfacial contact area plateaus is approximately equal to the theoretical limit for two hemispheres (0.33) at low *β*. In the high tension regime [Fig. 3e], we observe two adhesion regimes. Below a value of *A_M_* = 0.2, the interfacial contact area increases slowly with adhesion; beyond that value the interfacial contact area depends very weakly on adhesion.

Our doublet simulations demonstrate that our method is an accurate predictor of the mechanical interactions between cells. Furthermore, differences between simulation results and theoretical predictions further highlight the constraints of [Equation 2] in situations for which adhesion can’t be neglected. Importantly, our method doesn’t require the low adhesion assumption and can be used in any limit of adhesion or tension.

## 4. Quantitative predictions of sorting in MCAs

Having validated our method and used it to study cell doublet mechanics, we now demonstrate the influence of adhesion and interfacial tension on the sorting of two cell types in MCAs.

### 4.1. Methods to simulate cell sorting

In order to objectively to quantitatively assess the effect of mechanics on the extent and speed of self-organization in a cell aggregate, we developed three measures of sorting. These measures of sorting, as described below, are continuously calculated during the evolution of an MCA, and compared to a randomized reference distribution to produce a sorting index.

#### Radius measures

Since we focus here on spherical aggregates, two natural measures of the sorting are the mean distance of each cell type either from its own center of mass or from the center of mass of all cells. The former is a measure of cell type aggregation while the latter of sorting of one cell type to the boundary. In what follows, we focus on the mean radius of the cell type expected to aggregate from the center of mass of that same cell type, *X_r_*.

#### Neighbor measure

In a self-organizing system, we expect a higher probability of same-type cell neighbors. We consider two cells to be neighbors if at least one element in one cell interacts with an element of the other. It is via such interactions that neighbor cells apply forces to one another. The neighbor measure, *X_n_*, is then the like-like neighbor count for the cell type expected to sort to the center of the aggregate.

#### Surface measure

For spherical inside-outside sorting, we expect one cell type to occupy a greater proportion of the external surface of the MCA than the other. With our model, we are able to calculate the total area that is not part of a cell-cell interface across all cells. Therefore, for the surface measure we calculate the ratio, *X_s_*, of the external surface area occupied by the two cell types.

#### Comparison to randomized control distribution

To contextualize the measures described above, we introduced uniformly randomized control systems, against which the measures could be compared. To generate such controls, the fates of all cells in the system were randomly reassigned at each data output interval,while maintaining the same number of each cell type and the spatial distribution, at which point the sorting measures were then calculated again for these randomized systems [Fig. 4a]. Repeating this procedure 10000 times [Appendix B] produces a normal distribution of the sorting measures [Fig. 4b] from which we can calculate the mean, *E*(*X_i_*), maximum, *X_i,max,_* minimum, *X*_*i*,_*_*min*,_* and standard deviation, *σ*(*X_i_*), of each sorting measure *X*_*i*_. The measure as calculated from the simulation is then compared to the distribution of randomized measures by quantifying a sorting index, *S_i_* [Eq. 3]:

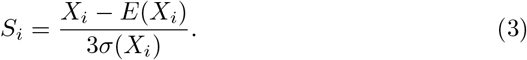

**Figure 4:**
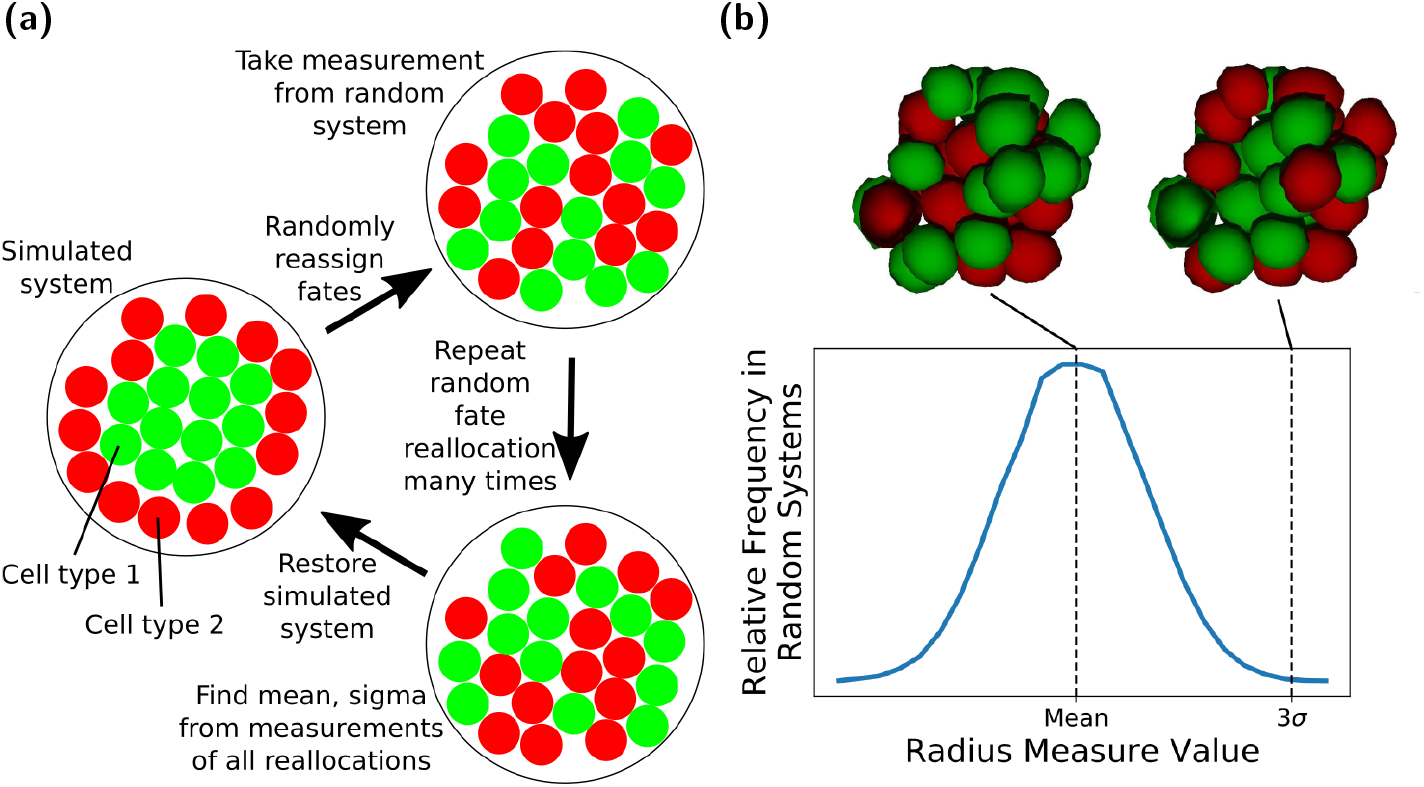
(a), Diagram showing how system fates are reassigned to produce mean, standard deviation, maximum, and minimum values of each measure for a given spatial configuration and cell type ratio. Starting from the simulation state, shown on the left, the fate of each cell is randomly reassigned, maintaining the same spatial distribution of cells, and the same ratio of cell types. All sorting measures are calculated for this new state. Fates are randomly reallocated up to 10000 times [Appendix B], to find the mean, maximum, minimum, and standard deviation of sorting measures, before returning the system to the original simulation state. (b), Distributions of radius sorting measure values over random fate reallocations for 10 and 30 cells. Distributions are seen to be approximately Gaussian, allowing calculation of mean, standard deviation, maximum, and minimum values. The same pattern is seen for other sorting indices. Spatial configurations at mean and 3*σ* shown.

The divisor serves as a way to ensure that highly deviant results in the randomized distribution don’t overly affect the sorting index.

For all following sorting simulations, we used our method to simulate MCAs growing from from 10 to 30 cells with two different cell types. We considered two cell types, cell type 1 and cell type 2. In the case of sorting, we define cell type 1 to be the type on the inside of the MCA [Fig.4a]. We calculated the radius, neighbor and surface sorting measures at regular intervals of each cell type, from which we quantified the respective sorting indices. We define 3 interfacial tension factors for like-like interfaces between both cell types (*β*_1,1_, *β*_2,2_) and unlike interfaces (*β*_1,2_). For simplicity in all simulations we set *β*_1,2_ = *β*_2,2_ = 1.0 and vary *β*_1,1_, henceforth simply referred to as *β*. Therefore, cell type 1 is by default the reference cell for calculating the sorting indices, i.e. the cell type that is assumed to aggregate on the inside of the MCA (see Figure 4). Unless otherwise stated, both cell types were given the same cortical tension which is justified given that Fig. 3 suggests that *γ_m_* is not a major driver of sorting. Furthermore, we restricted ourselves to the high cortical tension limit, and define the dimensionless adhesion parameter *α* = *A_M_/γ_m_* to simulate the dynamics of MCAs for a wide range of values of *α* and *β*. We also assume, unless otherwise stated, the same adhesion magnitude for both cell types, for both like and unlike adhesion. Experimentally, α is difficult to measure, but it is thought to be in the range of 0.2 to 0.25 at the most [13, 36, 12].

To test also the effect of a division bias, we ran two types of simulations for each pair of *α* and *β* values: (i) a symmetric division, in which in which each division produced cells of the same type as the parent; (ii) an asymmetric division, each dividing cell had a 50% change of producing two daughter cells of the same type as the parent, and 50% change of producing two daughter cells of different types.

All simulations were performed with both symmetric and asymmetric division, and each was run four times for each pair of α and β to account for variability in the simulations. The mean result and standard deviation for each set was calculated. The final value of sorting indices in simulations measures the extent of sorting in the final state of the system.

We will first use our method to test the hypothesis that doublet mechanics is a good predictor of sorting in an MCA, and then go on to use it to establish a better understanding of the mechanical drivers of cell sorting.

### 4.2. Doublet mechanics predicts sorting in MCAs

One of the main issues impeding the experimental validation of the mechanical drivers of sorting in MCAs is that it is very difficult to measure forces within an MCA. Therefore, cell doublets are often used to quantify the forces that can be used to predict sorting in an MCA. Therefore, we sought to test the hypothesis that cell doublets are a good and sufficient model for MCAs.

We ran the simulations for several random values *β* less than or equal to 1.0 and *α* less than or equal to 0.4 for both symmetric and asymmetric division. We quantified the sorting index for the final state of 30 different systems. We compared these values with the results of doublet testing [Fig. 3] with the same parameters. With this, we can assess how measurements taken from cell doublets predict the extent of sorting in MCAs. Fig. 5 shows final sorting indices plotted against the difference in doublet interface proportion measured for like-like doublets of two cell types with the same parameters as the corresponding MCA. The resulting scatter plots clearly show a strong positive correlation between the difference in interface proportion and the final sorting index across all sorting measures. This correlation is especially strong for systems with symmetric division. Importantly, this strong correlation establishes that mechanical doublet mechanics, which is experimentally tractable, can be used to predict the degree of sorting of MCAs.

**Figure 5:**
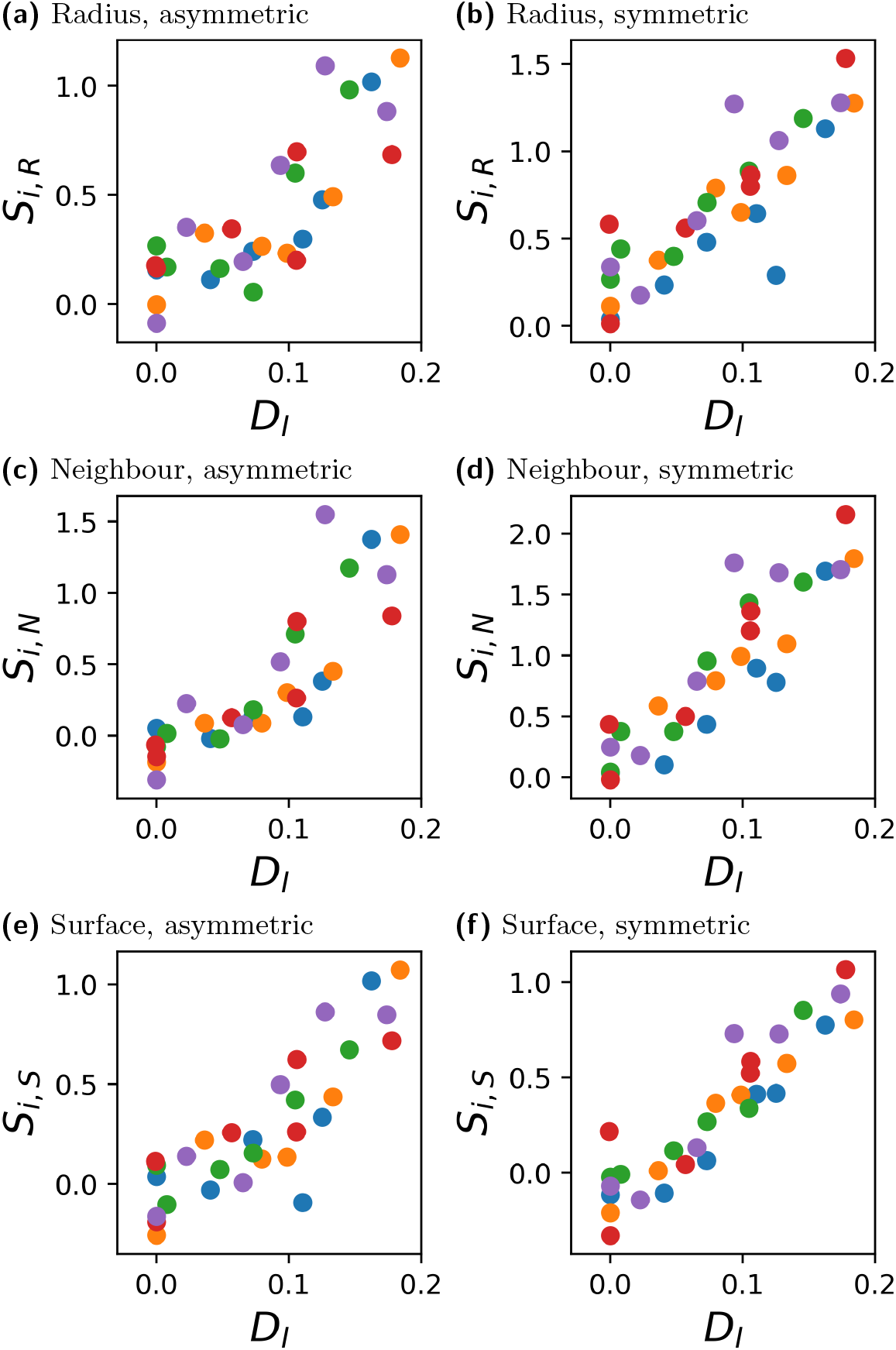
Plots of final sorting index for 30 cell aggregates against the difference in interface proportions for the two cell types found in doublet testing at corresponding parameter values. Each row of plots shows values of a different sorting measure for both symmetric and asymmetric division. A strong relationship is seen between the difference in interface proportion and final sorting index. Colour of point indicates adhesion magnitude.

### 4.3. Mechanical drivers of cell sorting in MCAs

In the following, we will use our simulations to make quantitative predictions about what adhesion and tension asymmetries are necessary for MCAs to sort. To do this, we performed simulations over a wide range of values of *α* and *β*. From the final value of sorting measures at the end of simulations we constructed a phase diagram in the *α* — *β* parameter space [Figs. 6a-6d]. From these phase diagrams we can deduce the limits on the values of α and *β* that drive sorting.

**Figure 6:**
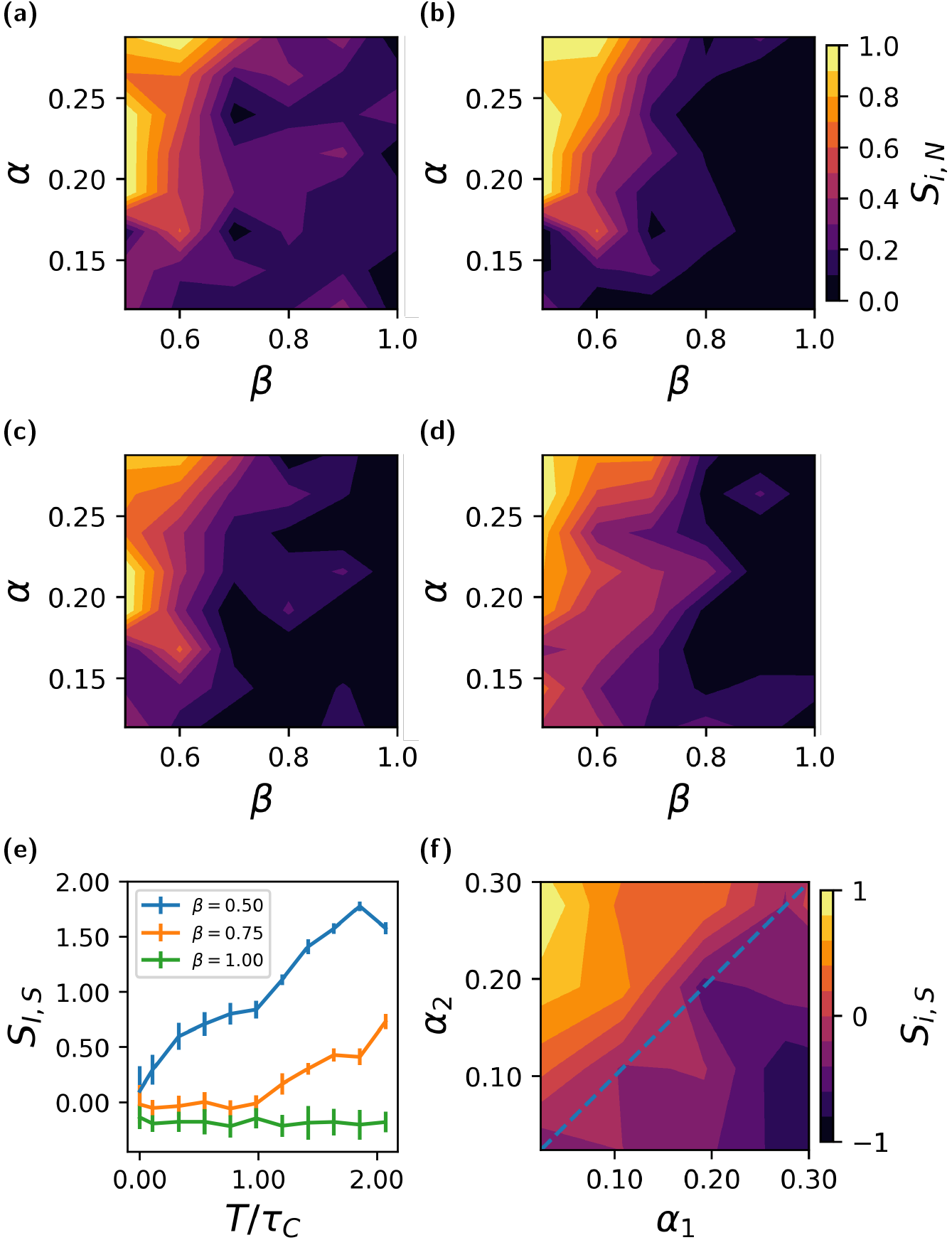
(a-d), Phase space plots of final sorting index for 30 cell systems in *α* and *β* space using (a), Radius measure, asymmetric division. (b), Neighbour measure, asymmetric division. (c), Surface measure, asymmetric division. (c), Surface measure, symmetric division. All cells have the same *γ_m_* and all cell types adhere with the same magnitude *α*. The tension at like-like interfaces for one cell type is altered by *β*. (e), Neighbour sorting index against *time/cell cycle time* in aggregates with asymmetric division and *α* = 0.2*γ_m_*. (f) Example of sorting by differential adhesion with asymmetric division from surface measure.

Figure 6a-6d demonstrate that there is good agreement between sorting measures about the behavior of the systems. Moreoever, the extent of sorting increases as *β* is reduced. This is predictable, given that a reduction of *β* is indicative of a relaxation of interfacial tension for cell type 1. Significantly, with our method we can now quantitatively predict the minimum of *β* ≈ 0.6 necessary to drive sorting at experimentally realistic values of *α*. We also see in Figure 6e that the kinetics of sorting are faster for smaller values of *β*, with very little sorting occurring for *β* = 0.75. It is also clear that a minimum adhesion is required in order for the *β* parameter to have any effect: for values of *α* < 0.2, little to no sorting is observed regardless of the value of *β*. This is interesting because *α* = 0.2 is in agreement with experimental observations of adhesion magnitude around 0.2 times cortical tension magnitude in systems where cortical tension has been proposed as a significant factor in morphogenesis [13, 12].

We also use our method to explore the effects of differential adhesion. As seen in [Fig. 6f] differential adhesion can drive sorting, but the difference in adhesion magnitude between the cell types has to be quite large - approximately a factor of 10 for full sorting. This suggests, in conjunction with the fact that adhesion forces are much smaller than tension forces, that tension asymmetry is a more essential factor in MCA sorting than adhesion asymmetry.

## 5. Discussion

We introduced a simulation method, which is an extension of SCEM, to investigate the mechanics of cell sorting. The model is based on local intra-and inter-cellular forces, and allows cell division and therefore evolution of any size MCA. It also allows us to define different regions of a cell and the mechanical properties of those regions. Therefore, our method is highly flexible and can model intercellular and intracellular mechanical interactions in a broad range of multicellular systems.

We first employed the experimentally tractable cell doublet system to both validate our numerical model and to probe the effect of adhesion and tension on the interfacial contact area between two cells. In a doublet, this contact area is determined by the balance of the adhesion force, cortical tension, and interfacial tension. We studied the effects of these parameters on the inner doublet’s interfacial contact area with our method and showed that it decreases sharply with interfacial tension before reaching a stationary phase in the high tension regime. We also showed that no interface can form when the adhesion drops below a threshold value. Above this value, the interfacial contact area depends only weakly on adhesion. Informed by experimental observations that tension forces are at least 4 times stronger than adhesion [13, 12], we studied the high-tension regime and found that the variation of interfacial contact area with the tension parameter, *β*, agrees remarkably well with the predictions of using local force balance at cell interfaces, neglecting adhesion. This validates our method, showing that it captures important aspects of the underlying mechanics.

Next, we simulated the evolution of MCAs, growing from 10 to 30 cells, to study the effects of adhesion and tension on the sorting process. We introduced three measures to quantify inside-outside sorting: (i) neighbour measure - the number of same-type nearest neighbours; (ii) radius measure - the mean spread of each type relative to its centre of mass; (iii) surface measure - the proportion of each cell type at the external surface of the aggregate. These measures were compared to their mean and standard deviation across a large number of random MCAs and used those to define corresponding sorting indices. The sorting indices we introduced are a novel and unbiased way to quantify sorting.

We observed a very strong correlation between the sorting indices in the MCAs and the interfacial contact area between cells in doublets. This observation is significant because, while experimental determination of mechanics within MCAs are difficult, measurements of cortical tension and contact angle in cell doublet are relatively straightforward. Furthermore, these plots [Figs. 5a-5e] suggest that relatively large differences in doublet interface area between the two cell types are required to produce complete sorting reliably.

Finally, in order to disentangle the relative effects of differential interfacial tension and differential adhesion, we constructed phase diagrams for the sorting indices as a function of *α* and *β*. These phase diagrams show that DIT alone is sufficient to produce full sorting of MCAs, as long as *β*, the relative cortical tension parameter, is lower than ≈ 0.6. Importantly, measurements of this experimentally accessible parameter in cell doublets can be used as a test of this prediction. Furthermore, we found that little to no sorting can take place when our dimensionless adhesion parameter fell below a value of *α* ≈ 0.2. This sets a lower bound on the adhesion level necessary to produce segregation in a tissue. Interestingly, this weak adhesion value is in agreement with the experimentally observed adhesion magnitude in the inner cell mass of the mouse embryo [13] and in the Drosophila eye [12]. Furthermore, the amount of differential adhesion necessary to exclusively drive cell sorting is comparatively large ≈ 10. This suggests that tension asymmetries are the primary driver of cell sorting.

Our results lead us to conclude that, for relatively small MCAs, similar to the sizes of those seen in early development, sorting can occur only if there are relatively large differences in either interfacial tension, adhesion or both between the different cell types. This could have vast implications in, for example, the early developing embryo where only few cells are present in an MCA.

Our method can be extended in several ways. For example, one can endow intra-cellular regions with different properties, making it possible to model the nucleus and the plasma membrane. Even more generally, this method can be extended straightforwardly to test aggregates that are not confined to roughly spherical geometries, many of which occur in biological organisms. For these reasons, our method can be useful well beyond studying tissue morphogenesis and could be applied to investigate a range of biological processes and pathologies, such as cancer cell metastasis.

## 6. Acknowledgments

The authors thank Tim Newman for providing the basic SCEM code for updating, and Ewa Paluch for valuable advice and input.

This work was performed using the Darwin Supercomputer of the University of Cambridge High Performance Computing Service (http://www.hpc.cam.ac.uk/), provided by Dell Inc. using Strategic Research Infrastructure Funding from the Higher Education Funding Council for England and funding from the Science and Technology Facilities Council.

## Appendix A. Decoupling Tension From Adhesion

**Figure A.7:**
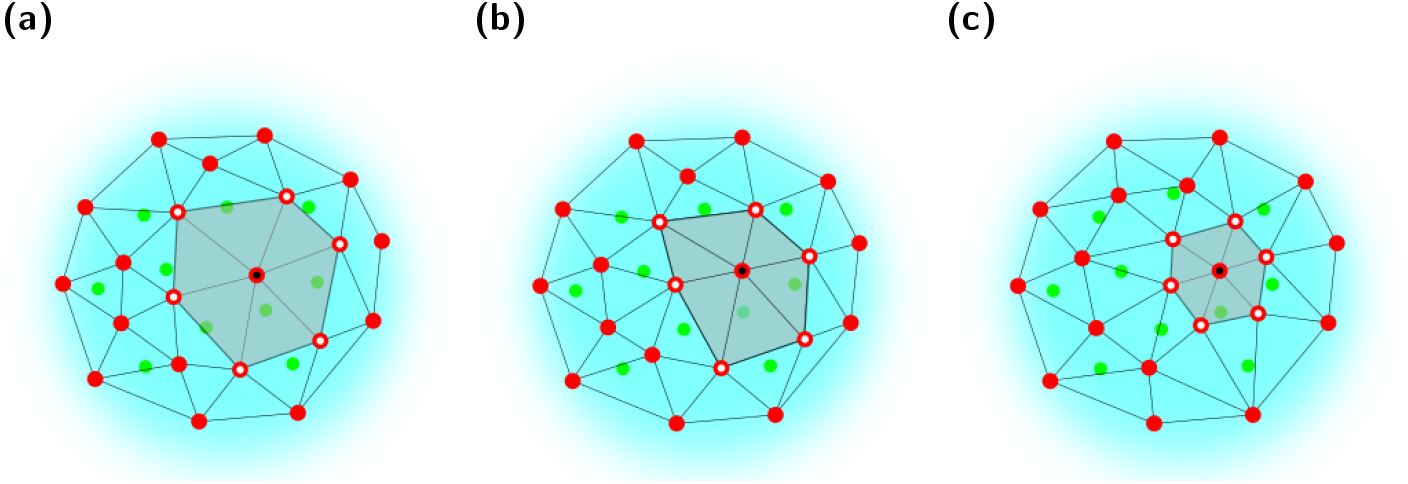
Diagram demonstrating how local area around an element is calculated to produce a normalisation factor that decouples the local adhesion magnitude from the local element density, and hence from the local cortical tension magnitude. The intercell adhesion magnitude of the element labelled with a black dot is multiplied by the grey-shaded area defined by all Delaunay triangles that have the element with a black dot as a vertex, divided by the mean area found for isolated cells in equilibrium. Thus the magnitude of adhesion is inversely proportional to the local density of elements. a, Interface over elements labelled with white dots has reduced interfacial tension. b, Interface over elements labelled with white dots has baseline tension. c, Interface over elements labelled with white dots has increased interfacial tension.

Care is required when implementing the DIT algorithm described in this paper. Changes to the local cortical tension magnitude in cortex-cortex interaction pairs will change the distance between elements in the pairs and thus result in changes to the local density of elements. Since adhesion between cells is mediated by these elements, a change in the element density will affect the local adhesion strength between neighbouring cells. This could counteract the expected effects of differential interfacial tension. For example, increasing the local tension at an interface, which should reduce the affinity between two cells, will result in a higher density of elements and thus a stronger local adhesion between the cells, potentially increasing their affinity in opposition to the effect of the change in tension.

To solve this problem, we devised an algorithm to normalise the adhesion magnitude of an element by the local element density. We begin by calculating the total area of all triangles in the Delaunay triangulation of cortex elements that have the element under consideration as one of their vertices [Figure A.7]. We then divide this area by the mean area for elements in a single cell at equilibrium to find a factor by which the area has changed relative to equilibrium. This factor is then used to change the magnitude of the adhesive interactions of the element. Thus any change from the equilibrium area will produce a corresponding change in local adhesion magnitude.

**Figure B.8:**
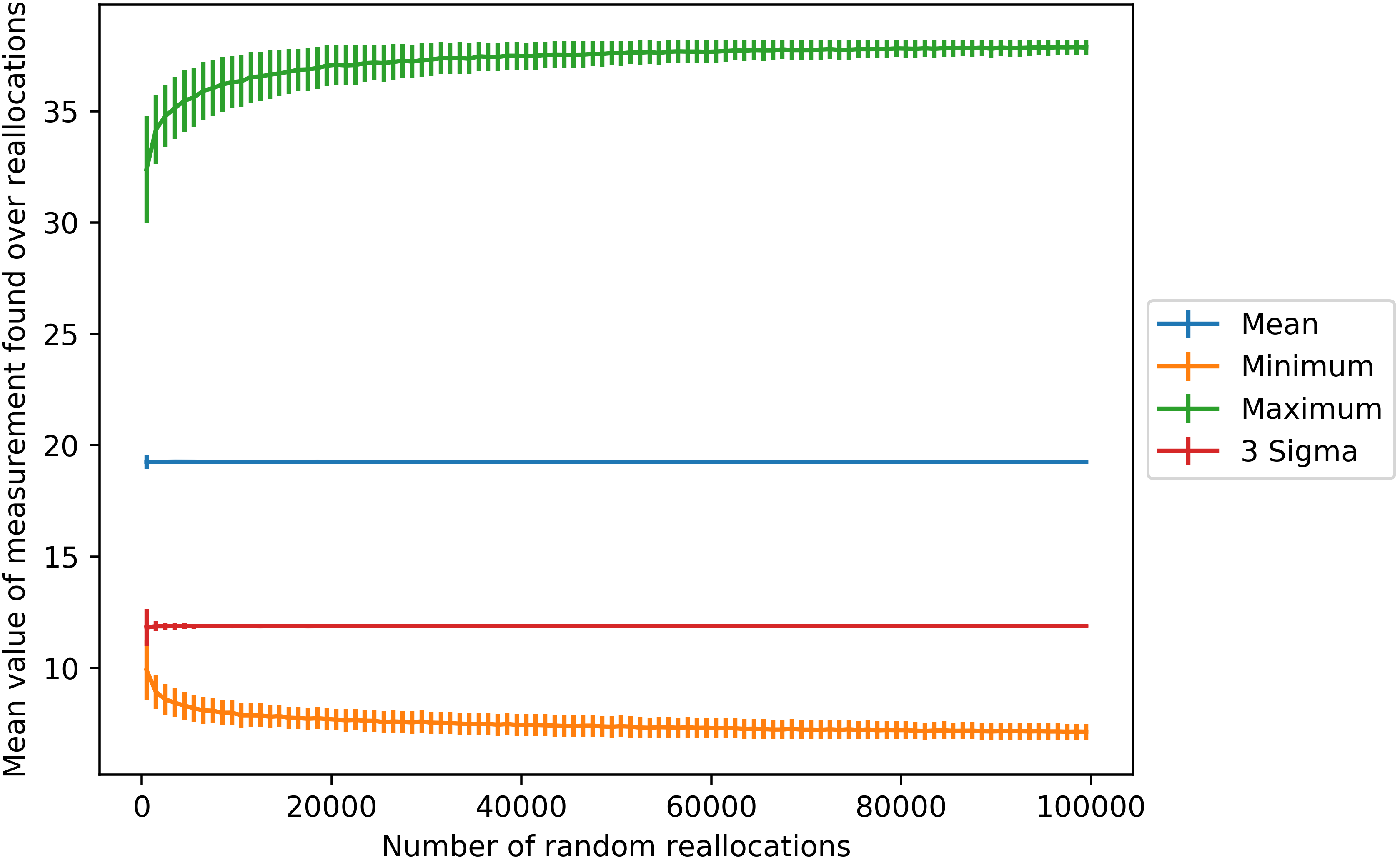
Variation of mean, maximum, minimum, and standard deviation in neighbour measure for a typical 30 cell system found over randomised tests against the number of random orientations tested. It can be seen that the values of these parameters are found with small errors for many fewer random tests than the total possible number of reallocations.

## Appendix B. Randomised Measure Distributions

The distribution distribution of values from randomised sorting measures is approximately Gaussian, allowing us to calculate a mean, standard deviation, maximum, and minimum value. Before performing the randomised measurement routine it’s important to establish how many random reallocations to perform. A 30 cell system with 15 cells of each type has almost 1.6 × 10^8^ possible different combinations of fates, and it is impossible to sample all such systems without increasing the run time of a simulation beyond what is computationally feasible. Fortunately, we were able to show that values quickly tend towards a limit over a much smaller number of repetitions. Figure B.8 shows how the mean, standard deviation, maximum, and minimum values found across random reallocations vary with the number of random reallocations tested for typical systems of 30 cells. It can be seen that the mean and standard deviation values are found with very little error over fairly small numbers of reallocations, whilst the maximum and minimum values have roughly found their limits after about 10^4^ reallocations. Thus we chose to use 10^4^ reallocations in our simulations. The exception to this is for small systems in which the total number of possible orientations is smaller than 10^4^, in which case the maximum is used instead.

